# Exploiting effector - host interactions using TurboID-based proximity labeling

**DOI:** 10.1101/2022.12.19.521009

**Authors:** Wei Shi, Sara C Stolze, Hirofumi Nakagami, Johana C. Misas Villamil, Isabel ML Saur, Gunther Doehlemann

## Abstract

Plant pathogens secrete effectors, which target host proteins to facilitate infection. The *Ustilago maydis* effector UmSee1 is required for tumor formation in the leaf during infection of maize. UmSee1 interacts with maize SGT1 and blocks its phosphorylation *in-vivo*. In the absence of UmSee1, *U. maydis* cannot trigger tumor formation in the bundle sheath. However, it remains unclear which host processes are manipulated by UmSee1 and the UmSee1-SGT1 interaction to cause the observed phenotype. Proximity-dependent protein labeling involving the turbo biotin ligase tag (TurboID) for proximal labeling of proteins is a powerful tool for identifying the protein interactome. We have generated transgenic *U. maydis* that secretes biotin ligase-fused See1 effector (UmSee1-TurboID-3HA) directly into maize cells. This approach, in combination with conventional co-immunoprecipitation allowed to identify additional UmSee1 interactors in maize cells. Collectively, our data identified three ubiquitin-proteasome pathway-related proteins (ZmSIP1, ZmSIP2, ZmSIP3) that either interact with or are close to UmSee1 during host infection of maize with *U. maydis.* ZmSIP3 represents a cell cycle regulator which degradation appears to be promoted in the presence of UmSee1. Our data provide a possible explanation for the requirement of UmSee1 in tumor formation during *U. maydis* - *Zea mays* interaction.

## Introduction

Most staple crops are essential for food security. Plant diseases greatly threaten food security by limiting the yield and quality of crops (Fisher *et al.*, 2016). Pathogens secrete effectors to target host structures for spreading infection (Dodds and Rathjen, 2010). Effectors usually modulate the biological activity of host proteins to suppress plant immunity (Hogenhout *et al.*, 2009). Identifying effector targets in the plant aids to the understanding of pathogenic mechanisms and thereby improves resistance to crop pathogens. The location, specificity, and low effector protein levels *in planta* are major obstacles for the identification of host target proteins.

*Ustilago maydis* is a plant fungal pathogen causing corn smut of many important crops and grasses, including the widely grown food and energy crop maize (*Zea mays)* (Kahmann and Kamper, 2004, Doehlemann *et al.*, 2008). *U. maydis* induces tumors on all aerial parts of maize and suppresses plant immunity by secreting effectors to the plant tissue (Skibbe *et al.*, 2010). Plant tumors produced upon *U. maydis* infection have been defined as clusters of cells that present abnormal cell division and reduced cell differentiation. The cell cycle is responsible for DNA replication and cell division and it is well-known that an abnormal cell cycle causes tumors in animals and plants (Jacobs *et al.*, 1999). *U. maydis* induces such tumors on all aerial parts of maize, which is a complex and dynamic process. The *U. maydis*-induced leaf tumor formation at the cellular level is composed of hypertrophic and hyperplasic tumor cells resulting from transformed mesophyll cells and bundle sheath cells, respectively (Matei *et al.*, 2018). About 460 effectors encoded by *U. maydis* target different host compartments (Lanver *et al.*, 2017). Only a few have been functionally characterized, including Pep1, Pit2, Cmu1, Tin2, See1, Mer1, Rip1 and Tips (Doehlemann *et al.*, 2009, Djamei *et al.*, 2011, Mueller *et al.*, 2013, Tanaka *et al.*, 2014, Redkar *et al.*, 2015a, Misas Villamil *et al.*, 2019, Darino *et al.*, 2021, Navarrete *et al.*, 2021, Bindics *et al.*, 2022, Saado *et al.*, 2022).

Interestingly, the organ-specific UmSee1 effector is required for the trans-differentiation of bundle sheath cells into hypertrophic tumor cells but not for the hypertrophic cells’ enlargement in the mesophyll cells, and the UmSee1-induced tumor formation and expansion only occur in vegetative tissues but not in floral tissues (Redkar *et al.*, 2015a). The UmSee1 effector is translocated from biotrophic hyphae to the plant cytoplasm and nucleus and a Yeast-two-Screen (Y2H) demonstrated the interaction of UmSee1 with maize SGT1 (suppressor of G2 allele of skp1) (Redkar *et al.*, 2015a). Several regulators control cell cycle progression, including cyclin-dependent kinases (CDKs), cell division control proteins (CDCs), cyclin proteins (CYCs), anaphase-promoting complex (APCs) and other activators or suppressors (Dang *et al.*, 2021). The previously identified UmSee1-interactor SGT1 is required for both the G1/S and G2/M cell cycle transitions in yeast and it is highly conserved in eukaryotic cells (Kitagawa *et al.*, 1999). Although, UmSee1-mediated inhibition of MAPK-triggered SGT1 phosphorylation could be detected (Redkar *et al.*, 2015a), it remains elusive, how SGT1 affects the plant cell cycle and how the cell cycle is influenced by UmSee1.

To give further insight into this key question and to minimize the individual limitations of each protein-protein interaction assay, we complement the previously performed Y2H screen (Redkar *et al.*, 2015a) with state-of-the-art *de novo* identification of UmSee1 interactors *in planta*. We apply standard co-immunoprecipitation (Co-IP) and establish proximity labeling (PL) for TurboID-catalyzed biotinylation of intracellular maize proteins upon *U. maydis*-mediated delivery of TurboID-fused UmSee1. Co-IP is very effective in obtaining stable complexes, but weak or transient interactors are not detected (Ngounou Wetie *et al.*, 2014, Sciuto *et al.*, 2019). PL captures weak or transient protein interactions and also proteins that locate consistently in close proximity of the protein of interest (Gingras *et al.*, 2019). Biotin-based PL approaches have been applied in yeast, plants and animals, including *Schizosaccharomyces pombe, Caenorhabditis elegans, Arabidopsis thaliana* and *Nicotiana benthamiana*, using transient expression or transgenic lines for specific cellular proteomes as well as interaction networks of nuclear, cytosolic and membrane baits (Branon *et al.*, 2018, Conlan *et al.*, 2018, Larochelle *et al.*, 2019, Mair *et al.*, 2019, Zhang *et al.*, 2019, Holzer *et al.*, 2022, Melkonian *et al.*, 2022). In the present study, we used a transgenic *U. maydis* strain to directly deliver UmSee1-TurboID-3HA fusion protein into maize cells during the infection process. This approach obtained distinct datasets of putative UmSee1-interacting candidates collectively suggesting a function of UmSee1 within the ubiquitin-proteasome network, a pathway also involving SGT1 (Dielen *et al.*, 2010). We selected three *<u>Z</u>ea <u>m</u>ays* Um<u>S</u>ee1 <u>i</u>nteracting <u>p</u>roteins (ZmSIPs) for further analysis and found them to also associate with SGT1. All three ZmSIPs have been previously implicated in the ubiquitin-proteasome pathway. Moreover, presence of UmSee1 is linked to a more rapid degradation of ZmSIP3, a cell cycle regulator 48 (CDC48), by the maize proteasome.

## Results

### Establishment of TurboID-based proximity labeling in the *U. maydis*-maize pathosystem

A previous study demonstrated that UmSee1 is secreted into host cells during *U. maydis* infection of maize leaves and that it can interact with the *Zea mays* SGT1 homolog (Redkar *et al.*, 2015a). To explore the potential for identifying further UmSee1-interacting proteins and proteins in close proximity to UmSee1 in maize leaves, we aimed to directly transfer UmSee-TurboID-3HA fusion proteins from biotrophic fungal hyphae into host cells (**Fig 1A**). We generated UmSee1 fused C-terminally to a TurboID-3HA tag and expressed it under control of the *pit2* promoter, which confers a high expression level in-planta (Doehlemann *et al.*, 2011). The construct (*pPit2:UmSee1-TurboID-3HA*, **Fig 1B**) was expressed in the *see1* knock out strain SG200Δsee1 strain which allowed to confirm that the recombinant effector protein fully complemented the virulence defect of the SG200Δsee1 strain (**Fig 1C, Table S3**). Anti-HA western blot analysis confirmed the successful enrichment of the fusion proteins (**Fig 1D**). To select specificity in the following experiments, the mCherry protein was fused to the signal peptide (SP) of UmSee1 (*pPit2:SP_UmSee1_-mCherry-TurboID-3HA*, **Fig 1B**) and expressed in *U. maydis* SG200, also under the *pit2* promoter. Maize leaves infected with the two *U. maydis* strains and treated with free biotin were subjected to protein extraction at three days post infection (dpi) under native (anti-HA) and denaturing (TurboID) conditions. Plant protein extracts were subjected to anti-HA immunoprecipitation and streptavidin-mediated isolation of biotinylated proteins, followed by liquid-chromatography mass spectrometry (LC-MS/MS) for the identification of isolated proteins (**Fig 1E**). Biotinylation of maize proteins by secreted UmSee1-TurboID-3HA was confirmed by strep-HRP western blot of raw protein extracts and streptavidin-enriched samples after treatment of the samples with 100 μM biotin (**Fig S1A**). Three biological replicates of Co-IP and TurboID were sampled from the identical plant for subsequent LC-MS/MS analysis. We detected enhanced biotinylation of maize proteins in all three replicates of samples carrying UmSee1 compared to the mCherry control samples (**Fig S1B**).

**Fig 1.:**
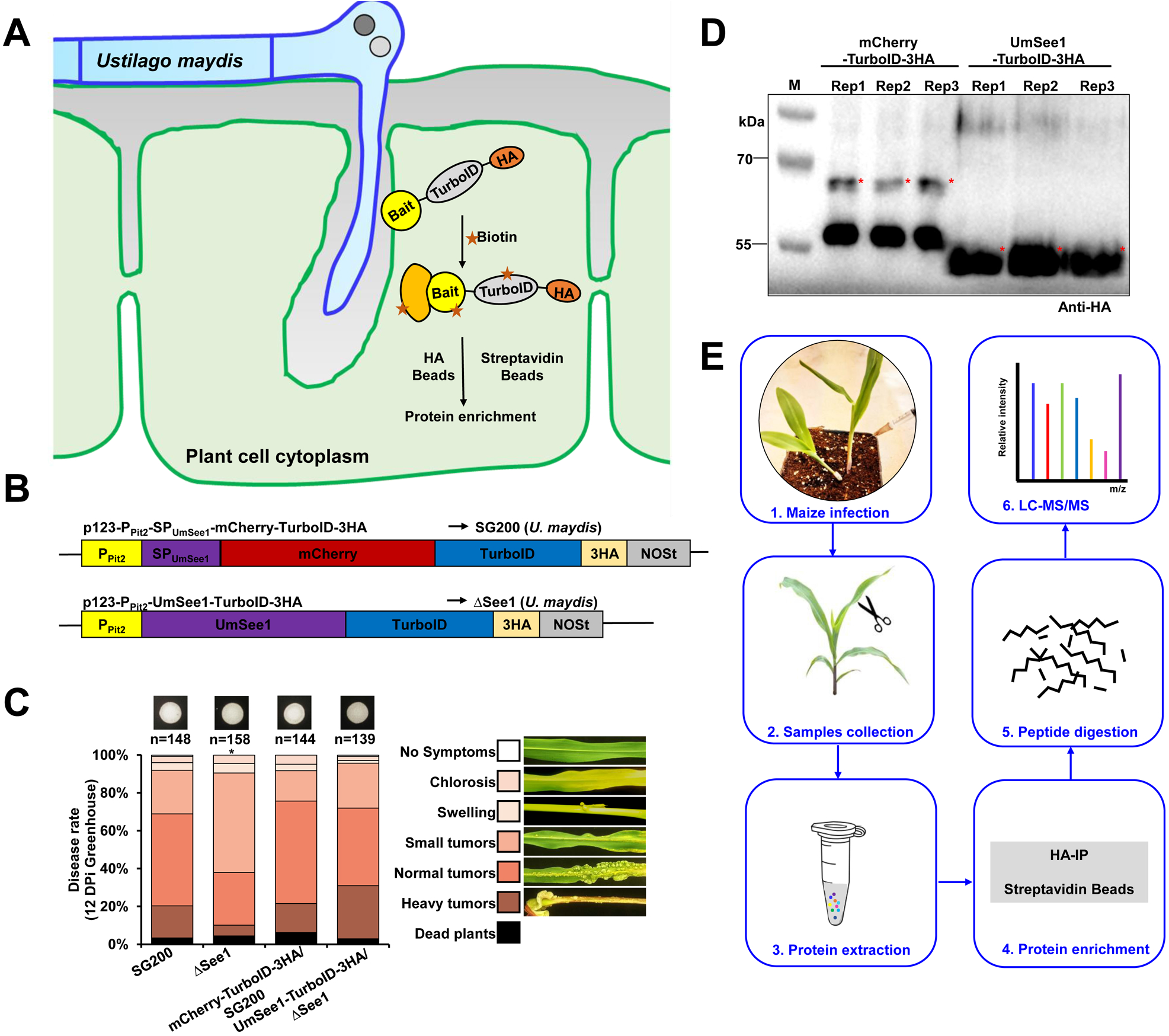
Experimental setup for TurboID-based biotin labeling in the *Ustilago-maize* system. (**A**) Principle to isolate further UmSee1-interacting proteins and proteins in close proximity to UmSee1 in maize leaves *vi*a Pull-down and TurboID-based proximity labelling (PL). (**B**) Schematic representation of the constructs used for HA-IP and Turbo-based PL. (**C**) Quantification of infection symptoms on EGB maize seedlings infected with *U. maydis* strains as indicated in B at 12 dpi. SG200: wild-type *U. maydis*, mCherry-TurboID-HA/SG200: mCherry-TurboID-3HA expressed in wild-type *U. maydis*, ΔSee1: *See1* deletion mutant, UmSee1-TurboID-3HA/ΔSee1: UmSee1-TurboID-3HA expressed in *See1* deletion mutant. (**D**) Detection of mCherry-TurboID-HA and UmSee1-TurboID-HA in maize leaves upon delivery by *U. maydis.* For anti-HA immunoprecipitation, leaves were harvested at three days post infection. Three independent replicates (Rep1-3) are shown. M: protein ladder. The asterisks represent target proteins. Expected size of mCherry-TurboID-3HA and UmSee1-TurboID-3HA proteins are 68.0 and 54.5 kDa, respectively. (**E**) Overview of the workflow used to identify putative UmSee1 targets by Co-Immunoprecipitation (Co-IP) and TurboID-based biotin labeling in the *Ustilago maydis*-maize pathosystem.

The relative amount of each identified protein within each replicate group by LC-MS/MS was quantified using label-free quantification (LFQ) and protein abundances with a sample were quantified using intensity-based absolute quantification (iBAQ) (Tyanova *et al.*, 2016). The data were analyzed by a t-test-based analysis (two valid LFQ values or more) and volcano plot analysis (three valid LFQ values). We identified 2563 and 1342 proteins in HA-IP and TurboID-based PL from the volcano plot analysis results, respectively (**Table S4**). High Pearson correlation values (*r* > 0.95) among the biological replicates of HA-IP and TurboID-based PL datasets indicate reproducibility of the data. To determine the interactors or enrichment, we used the RAW data to calculate two-sample t-tests with a permutation-based false discovery rate (FDR) of 5%. The enriched interactors of UmSee1 were designated if they had a p-value < 0.05 and log2 fold change > 1 over the control (**Fig 2A**). Finally, 262 and 67 proteins were identified by HA-IP and TurboID-based PL, respectively, and of these, 11 proteins were identified by both approaches (**Fig S1C, Table S5**). The datasets were further analyzed separately or together by psychophysiological interactions (PPIs) analysis, using the STRING database and GO (Gene Ontology) or KEGG (Kyoto Encyclopedia of Genes and Genomes) enrichment analysis using ShinyGO v0.66 (**Fig 2B-D, Fig S2, Fig S3**). PPIs and GO results suggest that the UmSee1 interactors and proteins in close proximity to UmSee1 *in planta* belong to the ubiquitin-proteasome complex.

**Fig 2.:**
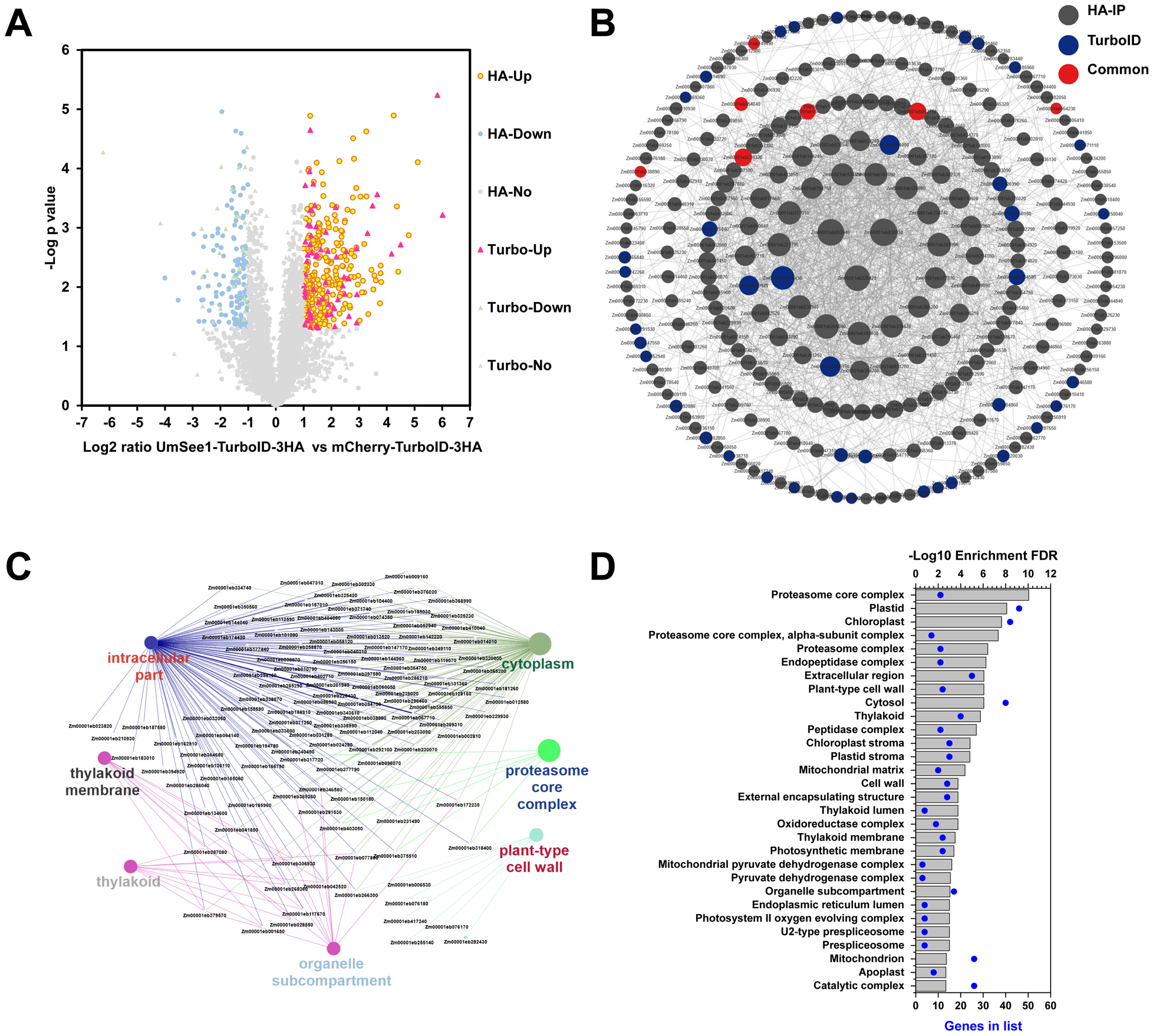
Identification of UmSee1 interacting proteins by co-immunoprecipitation (Co-IP) and TurboID-based biotin labeling. (**A**) Volcano plot analysis of identified proteins from HA-IP and TurboID-based biotin labeling by LC-MS/MS. Proteins significantly enriched in UmSee1 samples are shown in the top right corner (red and yellow dots). HA-up (yellow dots): significantly enriched proteins in UmSee1 compared to mCherry samples in the HA pull-down; HA-down (blue dots): significantly enriched proteins in mCherry compared to UmSee1 samples in the HA pull-down, HA-No (green dots): no significantly enriched proteins between UmSee1 and mCherry-samples in HA pull-down dataset, Turbo-up (red triangle): significantly enriched proteins in UmSee1 compared to mCherry samples in the TurboID-based proximity labeling; Turbo-down (purple triangles): significantly enriched proteins in mCherry compared to UmSee1 samples in the TurboID-based proximity labeling, Turbo-No (green triangles): no significantly enriched proteins between UmSee1 and mCherry samples in the TurboID-based proximity labeling dataset. (p-value < 0.05 and log2 fold change > 1 set as significantly enriched). (**B**) Protein-Protein interaction (PPI) analysis of the proteins identified in UmSee1 samples using the STRING database. Proteins significantly enriched in the HA-IP and TurboID-based PL of UmSee1 samples were submitted to the STRING database, and the protein interaction network was modified and analyzed using the Cytoscape software (version 3.9.1). Edges represent protein-protein associations, including known interactions and predicted interactions (gene neighbourhood, gene fusions, gene co-occurrence), the identified protein by the different methods were marked with different colors, and the shape’s size indicates the degree of interaction. Proteins are represented by their gene ID. (**C**) Cluster analysis of the Protein-Protein interactions network of the UmSee1-specific dataset by CluePedia (version 1.5.9) and ClueGo (version 2.5.9). (**D**) Gene Ontology (GO) cellular component enrichment of the UmSee1-specific dataset using ShinyGO v0.66.

### UmSee1 interacts with three UPS related proteins

Based on the analysis described above, our data suggest an involvement of UmSee1 with proteins of the ubiquitin-proteasome system. The previously identified UmSee1-interacting protein ZmSGT1 could be confirmed as an interactor in the HA-IP dataset. Importantly, SGT1 is a member of the SCF (Skp1/Cullin 1/F-box) complex, which associates with the ubiquitin-proteasome pathway (Cheng *et al.*, 2011). To determine the specificity and relevance of our approach to identified proteins of the ubiquitin-proteasome system, we chose three of the proteins identified by Co-IP and PL for further analysis. We named these three proteins ZmSIPs, for *Zea mays* See1-interacting protein. Zm00001eb157120 (ZmSIP1) was detected by anti-HA Co-IP and TurboID-based PL, but was not significantly enriched compared to the control. Zm00001eb369210 (ZmSIP2) was significantly enriched only in UmSee1 anti-HA Co-IP samples and Zm00001eb185960 (ZmSIP3) was significantly enriched only in TurboID-based PL datasets. Like ZmSIP2 and ZmSIP3, ZmSIP1 is also highly expressed in mesophyll cells but not in the bundle sheath cells, as demonstrated by cell-type specific transcriptome of maize leaves (Villajuana-Bonequi *et al.*, 2019) (**Fig S4**). Furthermore, the expression of ZmSIP2 and ZmSIP3 is affected by UmSee1 (Villajuana-Bonequi *et al.*, 2019), which might suggest that these genes could be involved in *U. maydis*-mediated tumor formation.

UmSee1 is localized in the nucleus and cytoplasm of plant cells (Redkar *et al.*, 2015a), and we therefore tested a putative subcellular co-localization of the ZmSIPs with UmSee1. The ZmSIPs were co-expressed as GFP function proteins in *N. benthamiana* leaves with UmSee1-mCherry or mCherry as the control. All genes were expressed under the control of the cauliflower mosaic virus (CaMV) 35S promoter. As expected, fluorescence microscopy detected the mCherry signal for UmSee1 in the nucleus and the cytoplasm. Similarly, GFP signals could also be detected in the nucleus and the cytoplasm (**Fig S5**), suggesting that all ZmSIPs (ZmSIP1, ZmSIP2, ZmSIP3) co-localize with UmSee1 in the same compartments upon co-overexpression *in planta.* To test for a direct protein-protein interaction between UmSee1 and the ZmSIPs, a directed yeast two-hybrid (Y2H) assay was done. The yeast transformants co-expressing each of the three ZmSIP1 proteins fused N-terminally to the GAL4 activation domain (pGBKT7 vector) with UmSee1, or mCherry fused N-terminally to the ADH1 activation domain (pGADT7 vector)) were selected on SD-Leu-Trp solid media and growth in the absence of histidine (-His), which indicates physical interaction of the proteins tested. In contrast to the BD-UmSee1/AD-mCherry transformants, the transformants expressing either of the AD-ZmSIPs together with BD-UmSee1 at OD600 < 1 (**Fig 3A**). The TurboID-based PL candidate interactor ZmSIP3 consistently appeared as the interactor with the lowest affinity to UmSee1 in the Y2H assays (**Fig 3A**). Importantly, we obtained comparable results when UmSee1 was exchanged to its ortholog from the maize-infecting smut species *Sporisorium reilianum*, SrSee1, in these assays (**Fig. S6**). In turn, when UmSee1 was exchanged to UhSee1, the ortholog from barley-infecting *Ustilago hordei*, we could not detect any growth in the absence of histidine (**Fig. S6**). This is in line with the previous finding that SrSee1, but not UhSee1 can restore the tumor formation of the UmSee1 deletion mutant (Redkar *et al.*, 2015b). We also analyzed protein-protein interactions by directed Co-IP assays. UmSee1ΔSP-6HA was co-immunoprecipitated by anti-myc immunoprecipitation of ZmSIP1-4myc, ZmSIP2-4myc, and ZmSIP3-4myc but not with GFP-4myc, suggesting that UmSee1 interacts with ZmSIP1, ZmSIP2 and ZmSIP3 *in planta* (**Fig 3B**). Again, the association of ZmSIP3 and UmSee1 appeared to be the weakest, although ZmSIP2 and ZmSIP3 protein levels were comparable in these assays. Similarly, in split-luciferase complementation imaging assays, a luminescence signal was detected when UmSee1 and all ZmSIPs were co-expressed in *N. benthamiana* leaves. We did not detect any signal in the control samples (**Fig 3C**). Collectively, our data shows that UmSee1 interacts with ZmSIP1, ZmSIP2 and ZmSIP3 and that the association with ZmSIP3 may be weak and/or transient interaction.

**Fig 3.:**
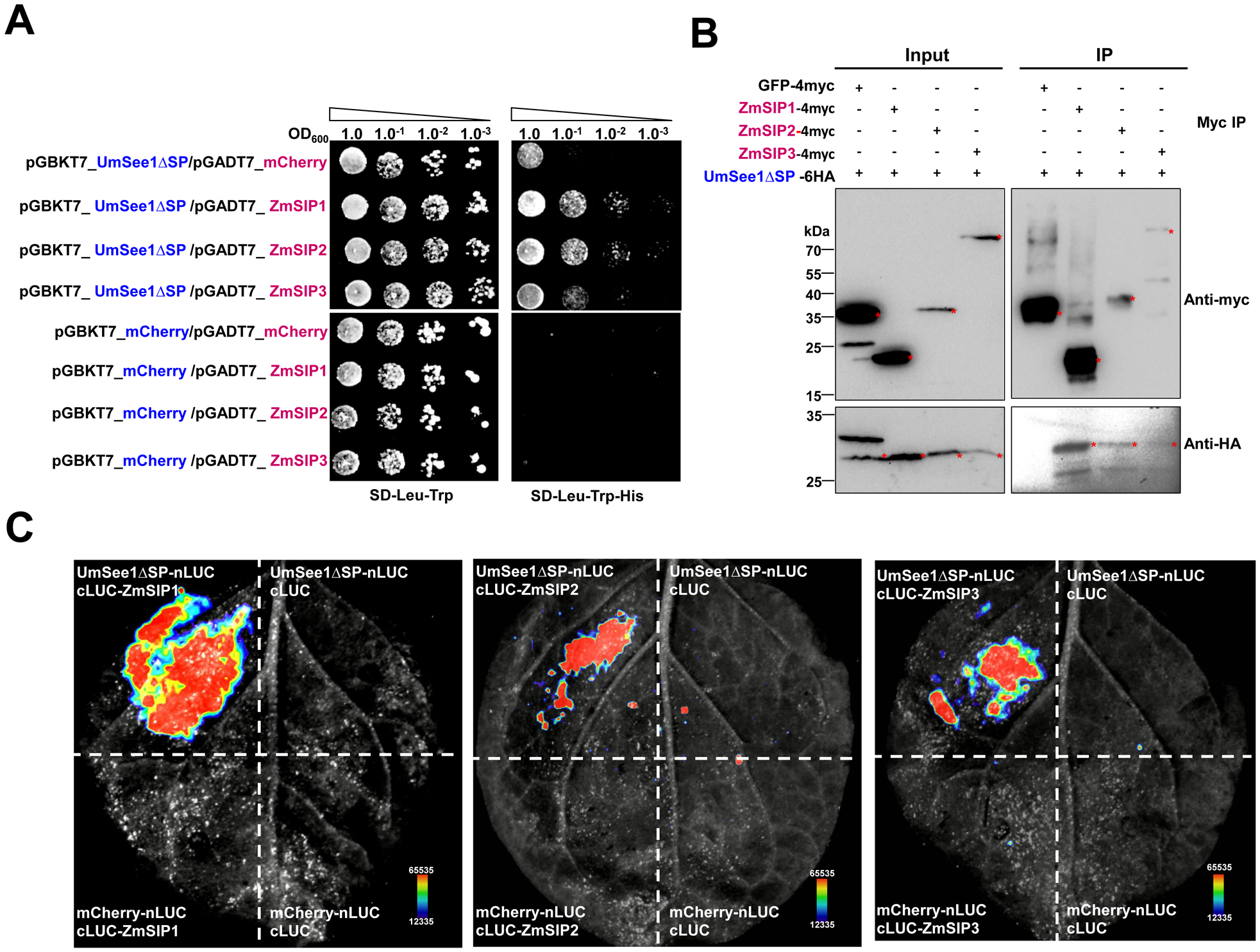
Three identified proteins enriched in theUmSee1 dataset interact with UmSee1 in one-to-one interaction assays. (**A**) Y2H assay of UmSee1 and three putative UmSee1 interaction partners (ZmSIPs). The yeast strain AH109 was co-transformed with mCherry or UmSee1ΔSP (UmSee1 without signal peptide) and the constructs of ZmSIPs (ZmSIP1, ZmSIP2, ZmSIP3). A drop-out assay of dilution series of the double-transformants was made in the presence (SD-Trp-Leu) and absence (SD-Trp-Leu-His) of histidine. Yeast growth in the absence of histidine suggests the interaction of proteins. Pictures of three independent experiments were taken 5 days after plating (**B**) Co-Immunoprecipitation assays of UmSee1 and the three ZmSIPs. *N. benthamiana* leaves were transiently transformed with the constructs encoding UmSee1ΔSP-6HA and ZmSIPs-4myc or GFP-4Myc. Leaves were harvested three days post-Agrobacterium-mediated transformation for protein extraction. Anti-myc immunoprecipitation (IP) was performed and total extracts and IP proteins were detected by western blot analysis using anti-HA and anti-myc. Expected size of GFP-4myc, ZmSIP1-4myc, ZmSIP2-4myc, ZmSIP3-4myc and UmSee1ΔSP-6HA proteins are 31.9, 20.5, 32.2, 86.1 and 26.4 kDa, respectively. (**C**) Split-luciferase assay of UmSee1 and the three ZmSIPs. *N. benthamiana* plants were transiently transformed with constructs of UmSee1ΔSP-nLUC or mCherry-nLUC and cLUC-ZmSIPs (ZmSIP1, ZmSIP2, ZmSIP3) or cLUC as indicated. Leaves were harvested two days post Agrobacterium-mediated transformation and treated with 1 mM D-luciferin for 10 min in the dark. Shown are representative pictures of luminescence signals from three independent biological replicates. Images were detected by the CCD imaging system (ChemiDoc, Bio-RAD).

### UmSee1-interactor maize proteins are components of the ZmSGT1-complex

SGT1 is known to regulate Skp1 in the SCF (Skp1/Cullin 1/F-box) complex, which controls the degradation of cell cycle regulators (Hermand, 2006). UmSee1 interacts with maize ZmSGT1 and can block SIPK-triggered SGT1 phosphorylation *(Redkar et al., 2015a)*. To explore the relationship between ZmSIPs (ZmSIP1, ZmSIP2, ZmSIP3) and ZmSGT1, we tested if ZmSGT1 can also associate with the here identified ZmSIPs. Yeast transformants expressing BD-ZmSGT1 with AD-ZmSIP1 and AD-ZmSIP2 grew on the selection medium lacking histidine. Transformants co-expressing BD-ZmSGT1 and AD-ZmSIP3 grew poorly in comparison, suggesting a stable association between UmSGT1 and ZmSIP1 or ZmSIP2, but a weak or transient association with ZmSIP3 (**Fig 4A**). ZmSGT1-6HA was also co-expressed with ZmSIP1-4myc, ZmSIP2-4myc, ZmSIP3-4myc or GFP-4myc in *N. benthamiana* leaves for Co-IP assays. ZmSGT1-6HA was precipitated at equal levels by anti-myc immunoprecipitation of ZmSIP1-4myc, ZmSIP2-4myc, and ZmSIP3-4myc but not with the GFP-4myc control (**Fig 4B**), suggesting that ZmSGT1 can interplay with ZmSIP1, ZmSIP2, and ZmSIP3 *in planta.*

**Fig 4.:**
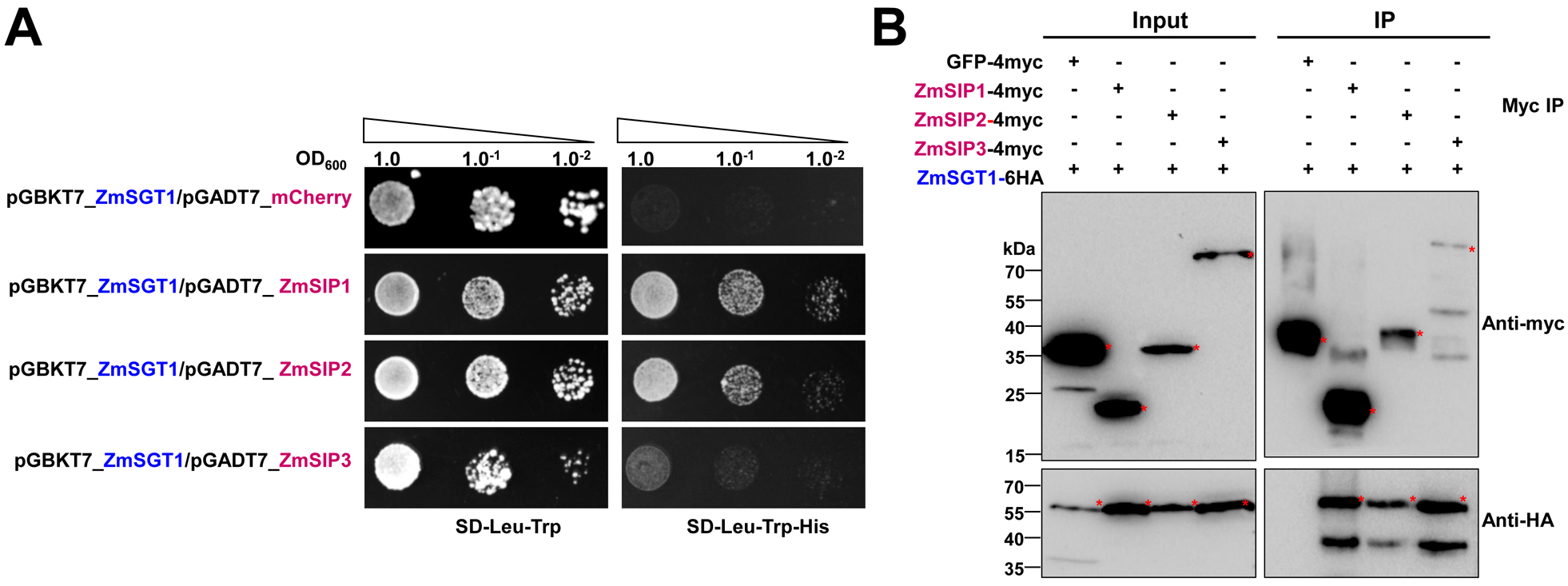
Three identified UmSee1-interactors are associated with the ZmSGT1-interacting protein complex. (**A**) Y2H assays of SGT1 and the three ZmSIPs. The yeast strain AH109 was co-transformed with ZmSGT1 and the three ZmSIPs, ZmSIP1, ZmSIP2, ZmSIP3, or mCherry as a negative control. A drop-out assay of dilution series of the double-transformants was made in the presence (SD-Trp-Leu) and absence (SD-Trp-Leu-His) of histidine. Yeast growth in the absence of histidine suggests the interaction of proteins. Shown are representative pictures of three independent biological experiments taken 5 days after drop out plating. (**B**) Co-Immunoprecipitation assays of ZmSGT1 and the ZmSIPs. *N. benthamiana* leaves were transiently transformed with the ZmSGT1-6HA and ZmSIPs (ZmSIP1, ZmSIP2, ZmSIP3)-4myc constructs. Leaves were harvested three days post-Agrobacterium-mediated transformation for protein extraction. Anti-myc immunoprecipitation (IP) was performed and total extracts and IP proteins were detected by western blot analysis using anti-HA and anti-myc. Expected size of GFP-4myc, ZmSIP1-4myc, ZmSIP2-4myc, ZmSIP3-4myc and ZmSGT1-6HA proteins are 31.9, 20.5, 32.2, 86.1 and 53.6 kDa, respectively.

### UmSee1 interacts with components of the UPS to regulate cell cycle

Since ZmSIP1, ZmSIP2 and ZmSIP3 are all proteins related to the ubiquitin-proteasome system, we performed an activity-based protein profiling (ABPP) assay to explore whether heterologous UmSee1 affects proteasome activity. For this, total proteins extracted from maize leaves were mixed with or without recombinant 6His-UmSee1 obtained from *E. coli*. Samples were pre-incubated in the presence or absence of the proteasome inhibitor epoxomicin and labeled with the specific proteasome activity-based probe MVB072 (Kolodziejek *et al.*, 2011). MVB072 binds covalently and irreversible to the active site of the catalytic proteasome subunits beta-1, beta-2 and beta-5 and contains a bodipy tag for fluorescent detection and a biotin tag for affinity purification (Kolodziejek *et al.*, 2011). Proteasome specific signals were observed at ca. 26 kDa and those signals were inhibited in the presence of epoxomicin (**Fig 5A**). To compare, if in the presence of See1 proteasome activity is enhanced ABPP signals were quantified and percentage of activity compared to buffer was plotted. We detected significantly enhanced activity of the maize proteasome in the presence of recombinant 6His-UmSee1ΔSP in comparison to the control sample. (**Fig 5A-B**). Thus, addition of recombinant 6His-UmSee1 enhances *in-vitro* proteasome activity of maize. To explore the effect of UmSee1 on the ubiquitination of maize proteins, seven-day-old maize leaves were infected with *U. maydis* strains SG200 or SG200ΔSee1. Total extracted proteins were detected via western blot using an α-ubiquitin antibody. We consistently detected stronger ubiquitination in SG200-infected leaves when compared to SG200ΔSee1-infected leaves, suggesting that UmSee1, directly or indirectly, increases the ubiquitination of maize proteins during infection (**Fig 5C**). Interestingly, the expression pattern of the proteasome alpha subunit ZmSIP2 is downregulated in SG200ΔSee1-infected leaves in comparison to SG200 infection but the expression of the catalytic beta-1 proteasome subunit PBA1 (GRMZM2G1775) does not change during infection (Fig S4) (Villajuana-Bonequi *et al.*, 2019), suggesting a normal core protease proteasome activity but a somehow disrupted structure of the 20S barrel in the absence of UmSee1. How could one then connect UmSee1-dependent proteasomal activity to its role in the activation of plant cell cycle? A possible link could be ZmSIP3, which contains CDC48 and AAA+ domains related to cell cycle regulation. The protein stability of ZmSIP3 may be important for cell cycle. To explore if the UmSee1-driven effects on proteasome-activity and ubiquitination alter the stability of ZmSIP3, we performed a protein degradation analysis of ZmSIP3 in the presence of UmSee1ΔSP-mCherry-6HA, as well as an mCherry-6HA control (Fig 5D). For this, ZmSIP3-4myc, mCherry-6HA and UmSee1 ΔSP-mCherry-6HA were separately expressed in *N. benthamiana* leaves using Agrobacterium-mediated transformation. ZmSIP3-4myc protein extracts were supplemented with increasing concentrations of mCherry-6HA or UmSee1ΔSP-mCherry-6HA-containig extracts in the presence or absence of the proteasome inhibitor MG132. Samples were incubated at 28°C for 45 min, and the amount of ZmSIP3-4myc was detected by western blot using anti-myc antibodies. ZmSIP3-4myc protein levels decreased with increasing UmSee1ΔSP-mCherry-6HA concentrations and this degradation process was reversed by the addition of MG132. This decrease of ZmSIP3-4myc was significantly less seen when UmSee1ΔSP-mCherry-6HA was exchanged by mCherry-6HA. These data indicate an enhanced proteasome-mediated degradation of ZmSIP3 in the presence of UmSee1. Collectively, our results suggest a possible UPS regulatory network modulated by UmSee1 that facilitates cell cycle progression and thus promotes tumorigenesis (**Figure 5E**).

**Fig 5.:**
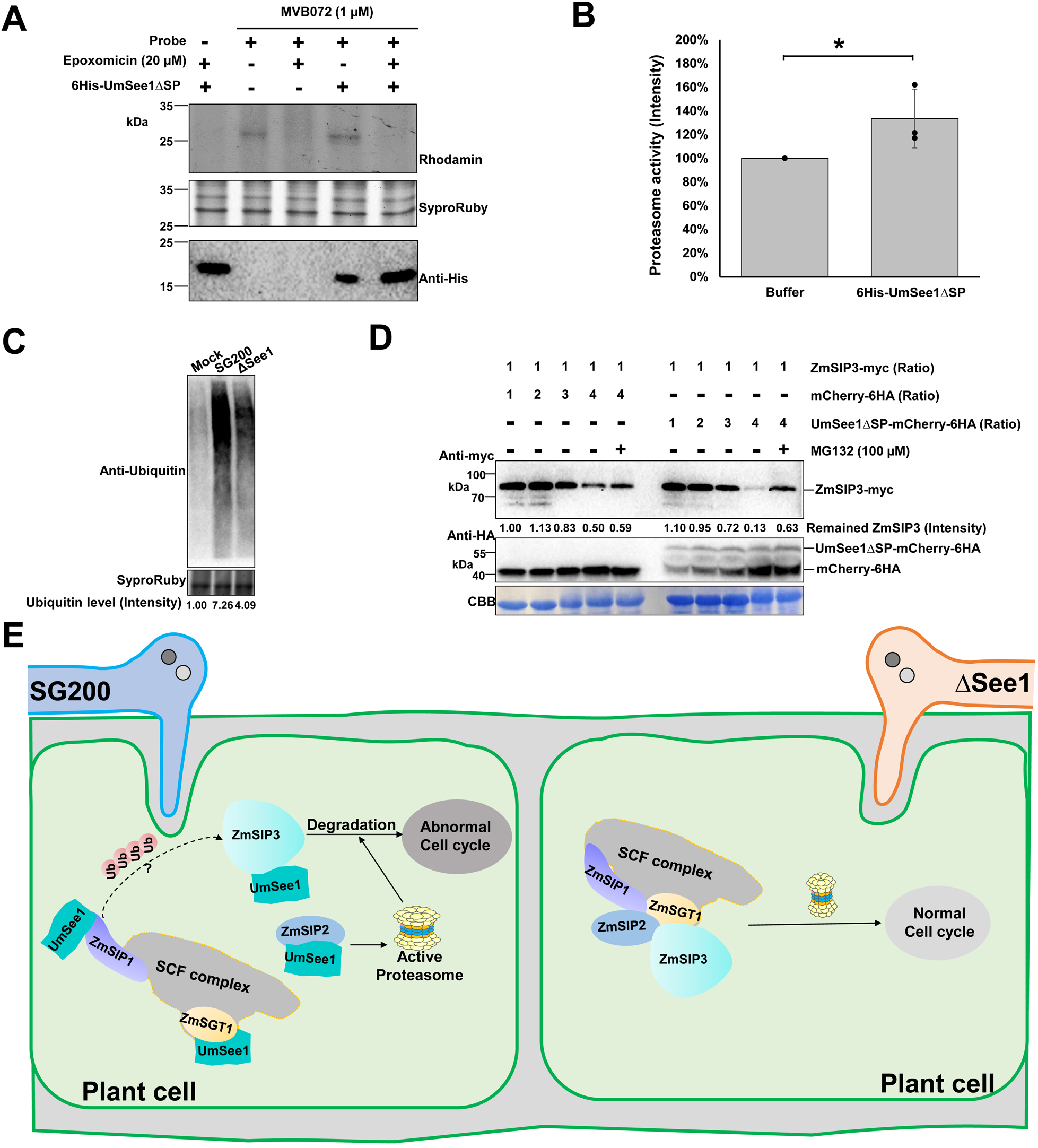
UmSee1 enhances proteasome activity and can promote ZmSIP3 degradation. (**A**) Activity-based protein profiling (ABPP) of the effect of recombinant UmSee1 on proteasome activity. Total proteins extracted from maize leaves were mixed with the probe MVB072. Samples were pre-incubated with or without recombinant UmSee1_His and with or without the proteasome inhibitor epoxomicin. Samples were analyzed by gel electrophoresis and monitored using a Rhodamine filter. SyproRuby staining served as loading control. Purified 6His-UmSee1ΔSP were detected by western blot using an anti-His antibody. (**B**) Relative proteasome activity quantification of (A). (**C**) Maize ubiquitin activity upon infection with *U. maydis* strains SG200 and SG200ΔSee1. Leaves were harvested three days post-infection and total protein extracts were analyzed by anti-ubiquitin western blot. SyproRuby staining serves as loading control. (**D**) In vitro protein degradation analysis of ZmSIP3. ZmSIP3-4myc, mCherry-6HA and UmSee1 ΔSP-mCherry-6HA were separately expressed in *N. benthamiana* leaves. Total ZmSIP3-4myc protein lacking protease inhibitor were mixed in different proportions with mCherry-6HA or UmSee1ΔSP-mCherry-6HA protein extracts either in the presence of 100 μM MG132 or DMSO (control) and incubated at 28°C for 45 min before the reactions were stopped by boiling samples at 95°C in 2xSDS loading buffer for 10 min. Samples were analyzed by western blot using anti-myc and anti-HA as indicated. Coomassie brilliant blue (CBB) staining served as loading control. (**E**) Hypothetical model for the function of UmSee1. *U. maydis* secretes UmSee1 protein which binds to ZmSGT1 and ZmSIPs (ZmSIP1, ZmSIP2, ZmSIP3) to modify the SCF complex and enhance proteasome activity, affecting the degradation of cell cycle-related proteins, including ZmSIP3.

## Discussion

### Combination of AP-MS and PL identifies the UmSee1-interaction network

Plant pathogens target host proteins with secreted effectors to facilitate infection. Thus, the identification of effector targets helps to understand both, microbial virulence strategies, as well as mechanisms of plant immunity. Conventional approaches for identifying protein interactors are yeast-two-hybrid (Y2H) library screenings and affinity purification followed by mass spectrometry (AP-MS) (Paraoan *et al.*, 2020, Woloschuk *et al.*, 2020). ZmSGT1 was identified as an interactor of UmSee1 via a yeast-two hybrid library screen and this interaction was confirmed by AP-MS (Redkar *et al.*, 2015a). However, this strategy likely fails to explore complex effector-target interaction networks, which can be dynamic and transient in nature. Plant pathogen effectors can target multiple host proteins to effectively disrupt a single or various biological pathways and promote virulence (Khan *et al.*, 2018). For example, *Pseudomonas syringae* effector HopM1 (HopPtoM) is required for full virulence by targeting MIN7 (ARF-GEF), MIN10 (GRF10), UPL1, UPL3, and ECM29 proteins in Arabidopsis (Toruño *et al.*, 2016, Üstün *et al.*, 2016, Khan *et al.*, 2018). Another example is the effector AvrPiZ-t from *Magnaporthe oryzae*, which targets Nup98 homolog APIP12 and the E3 ligase APIP6 and APIP10 in rice to promote infection (Park *et al.*, 2012, Park *et al.*, 2016, Tang *et al.*, 2017).

Enzyme-catalyzed proximity labeling approaches have been increasingly used as an alternative screening method for capturing weak or transient protein interactions by labeling the neighboring proteins with biotin (Gingras *et al.*, 2019). TurboID-based PL is ideal for low affinity, transient interactions, and proteins insoluble under native extraction conditions (Zhang *et al.*, 2019, Melkonian *et al.*, 2022). The components of regulators and many subcellular organelles involved in various cellular processes have been identified by TurboID-based PL in mammalian cells and some model plants, but it has not been yet used in the context of fungal pathogens.

In this study, we established TurboID-based PL for exploiting the so far unidentified interactors of the organ-specific effector UmSee1 in the *U. maydis*-maize pathosystem. Combination of all putative UmSee1 interacting proteins identified by co-immunoprecipitation (Co-IP) and TurboID-based PL identified components of the ubiquitin-proteasome system (UPS) being enriched in the UmSee1 interaction network. Three UPS-related proteins (ZmSIP1, ZmSIP2, ZmSIP3) were confirmed to interact with UmSee1. ZmSIP1 contains a ubiquitin-conjugating enzyme E2 motif, catalytic (UBCc) domain, suggesting that ZmSIP1 in maize may perform as a E2 ubiquitin-conjugating enzyme. E2 enzymes have been found to regulate the processing and topology of poly-ubiquitin chain formation and other modifications, thus, determining the destiny of the modified proteins (Zhou *et al.*, 2017). The interaction of UmSee1 and ZmSIP1 may be linked to the proteasome alpha subunit ZmSIP2 and the increased ubiquitination observed upon *U. maydis*-infection, but the molecular mechanism behind this phenotype needs to be further investigated. ZmSIP2 contains a proteasome alpha type 3 domain, which is part of the central core 20S proteasome containing the catalytic subunits responsible for protein degradation in the cytoplasm and nucleus. The plant proteasome is a key regulator of protein stability and has been identified as a hub in plant immunity since it is targeted by multiple bacterial and fungal effector molecules, such as SylA and HopZ4 from *Pseudomonas syringae* or higginsianin B from the fungus *Colletotrichum higginsianum* that bind and inhibit proteasome activity suppressing plant immunity (Misas-Villamil *et al.*, 2013, Üstün *et al.*, 2014, Dallery *et al.*, 2020).

### Towards an understanding of effector-mediated modulation of cell-cycle control

UmSee1 interacts with and alter the function of UPS components. The observed UmSee1-mediated increase in proteasome activity suggests that this effector affects proteasome function or regulation, although it is not yet clear if the *in vitro* proteasomal activation observed is specific for UmSee1 or if it is a consequence of UmSee1 degradation where UmSee1 is used as a substrate. If UmSee1 specifically activates proteasomal degradation, one would expect an increased degradation efficiency of certain proteins. In this context, it is important that essential cell cycle regulators are associated with the UPS, and that UPS regulates cell growth and proliferation boundaries between the G1 and S-phase (Bashir *et al.*, 2004). The UmSee1 interactor ZmSIP3 is a CDC48 domain-containing protein, i.e. a conserved AAA+ (ATPases Associated with diverse cellular Activities) ATPase, which are enzymes known to be involved in cell cycle control, cytokinesis and cell proliferation (Stolz *et al.*, 2011). CDC48 has been first identified in yeast, where it is important for cell cycle progression (Buchberger, 2013). In plants, CDC48 is involved in multiple processes, such as germination, reproduction, tissue differentiation and immune response. For example, *A. thaliana* CDC48A is involved in embryo growth and the degradation of immune receptors (Li *et al.*, 2022). Thus, the increase of UPS-mediated degradation of ZmSIP3 in presence of UmSee1 might provide a hint on how *U. maydis* triggers cell proliferation in host. Based on our findings, a possible scenario (**Fig 5E**) is that UmSee1 interferes with components of the SGT1/SCF complex to potentiate UPS activity. This leads to an increased degradation of cell cycle regulators, including CDC48, which promotes the activation of cell cycle progression observed in *U. maydis* induced tumorigenesis. In future studies, this working model can be tested to dissect the molecular interplay between the ubiquitin-proteasome system and the tumor-inducing function of UmSee1. In summary, we have established TurboID-based PL in combination with Co-IP followed by LC-MS/MS to identify a protein interaction network of the UmSee1 effector. This approach is widely applicable, in particular for establishing interaction networks of effectors which interfere with complex cellular processes. Insight into protein networks and their modulation by effectors will increase our mechanistic understanding of microbial pathogenesis. Moreover, it provides new starting points for the biochemical and structural elucidation of effector-target interactions and therefore holds a strong potential to establish new strategies for plant disease control and crop improvement.

## Materials and methods

### Plasmid construction

NEBuilder HiFi DNA Assembly (Gibson cloning method) was used for all cloning processes. For the transformation of *U. maydis*, the plasmid p123-PPit2-TurboID-3HA containing the native promoter of UmPit2 (UMAG_01375), TurboID and 3HA tag was generated using NEBuilder HiFi DNA Assembly. This plasmid was then used to generate the p123-PPit2-SPUmSee1-mCherry-TurboID-3HA and the p123-PPit2-UmSee1-TurboID-3HA also using NEBuilder HiFi DNA Assembly. All constructs were confirmed by sequencing. For Y2H assays, cDNAs of mCherry and ZmSIPs (ZmSIP1, ZmSIP2, ZmSIP3) were cloned into the pGADT7 vector (Clontech) for expression *via* the GAL4 AD under constitutive ADH1 promoter. mCherry, UmSee1ΔSP, SrSee1ΔSP, and UhSee1ΔSP were cloned into the pGBKT7 vector (Clontech) for expression with the GAL4 DNA-BD under control of the constitutive ADH1 promoter. For CoIP and *in vivo* protein degradation assays, cDNAs of GFP and ZmSIPs (ZmSIP1, ZmSIP2, ZmSIP3) lacking stop codons were cloned into pICH47732 vector with the cauliflower mosaic virus 35S promoter and a 4myc tag, CDSs of UmSee1ΔSP and ZmSGT1 were cloned into pICH47732 vector with the cauliflower mosaic virus 35S promoter and a 6HA tag. For subcellular localization studies, ZmSIPs (ZmSIP1, ZmSIP2, ZmSIP3) lacking stop codons were cloned into pICH47732 vector with the 35S promoter and a GFP tag. UmSee1ΔSP was cloned into pICH47732 vector with the 35S promoter and a mCherry tag. For UmSee1ΔSP protein expression in *E. coli* and subsequent ABPP, UmSee1 CDS was amplified by PCR without its signal peptide and ligated into the pET15b vector. All primers used are listed in **Table S1**.

### Generation of fungal strains and growth conditions

*U. maydis* strains generated and used in this study are listed in **Table S2**. p123-P_Pit2_-SPUmSee1-mCherry-TurboID-3HA by p123-P_Pit2_-TurboID-3HA The p123-P_Pit2_-UmSee1-mCherry-TurboID-3HA and p123-P_Pit2_-SP_UmSee1_-mCherry-TurboID-3HA constructs containing the carboxin resistance *ip* allele was introduced into SG200ΔSee1 and SG200, respectively, by homologous recombination in the *ip*-locus (Kämper, 2004) All generated *U. maydis* strains were confirmed by PCR and Southern blot.

### Maize infections and biotin-treatment

*U. maydis* strains were grown in liquid YEPS light medium on a shaker (200 rpm) or potato dextrose agar (PD) plates at 28 °C. For assessing virulence, the seven-day-old maize seedings of Early Golden Bantam (EGB) were inoculated with *U. maydis* strains with an OD_600_=1.0. Disease symptoms were scored at 12 days post-infection (dpi) as described (Kämper *et al.*, 2006) and the experiment was performed at least three times independently. For the preparation of samples for LC-MS/MS, seven-day-old maize seedings of Early Golden Bantam (EGB) were inoculated with respective *U. maydis* strains at an OD_600_=3.0. Biotin treatment of the maize leaves was performed via infiltration of a 100 μM biotin solution at two dpi.

### Sample preparation for mass spectrometry (MS)

For each construct (*pPit2:SP_UmSee1_-mCherry-TurboID-3HA,pPit2:UmSee1-TurboID-3HA*), three biological replicates were performed and analyzed via MS. The biotin-treated maize leaves were harvested at three days post-infection (2 days post biotin treatment) and frozen in liquid nitrogen. Frozen plant material was ground to a fine powder for protein extraction. These plant powders were divided into different tubes. For HA-IP, the frozen powder was added to 1.5 mL ice-cold protein-lysis buffer (50 mM Tris pH 7.5,150 mM NaCl,10 % glycerol, 2 mM EDTA, 10 mM DTT, 0.5 % IGEPAL, 1 mM PMSF) with cOmplete™ Protease Inhibitor Cocktail (Roche) and incubated on ice for half an hour after vortexing. The tubes were centrifuged at 13,300 x g for 10 min at 4°C, the upper soluble fraction was centrifuged again at 13,300 x g for 20 min at 4°C, the resulting supernatant was transferred to a fresh tube as total protein extract. HA magnetic beads (Pierce Anti-HA magnetic beads, Thermo Fisher Scientific) were separated on a magnetic separator. and washed three times by ice-cold wash buffer (50 mM Tris–HCl pH 7.5, 150 mM NaCl, 10% glycerol, 2 mM EDTA). After adding 10 μL beads to each of the cell extracts, the samples were incubated on a rolling wheel at 4°C for three hours. HA magnetic beads were magnetically separated and washed in 1 mL wash buffer three times. An aliquot of the beads was used for immunoblotting, the antibodies used were mouse anti-HA primary antibody (1:30,000 dilution; Sigma-Aldrich) and anti-mouse IgG secondary antibody (1:10,000 dilution; Thermo Fisher Scientific).

For TurboID-based PL, the frozen powder was added to 500 μl SDT-lysis buffer (10 mL 10% SDS, 2.5 ml 1M DTT, 2.5 mL 1M Tris pH 7.5, 10 mL ddH_2_O)(Melkonian *et al.*, 2022), and incubated 5 min at 95°C after vortex-mixing. The samples were sonicated for 10 min and centrifuged for 10min at 13, 300 x g. The resulting supernatant was transferred to a fresh tube as total proteins. Biotin depletion of the total proteins used the methanol-chloroform-precipitation method (Melkonian *et al.*, 2022). 666 μL methanol and 166 μL chloroform were added to 500 μL total protein extracts and mixed with 300 μL ddH_2_O. The mix was centrifuged for 10 min at 1,500 g and the supernatant was discarded, the protein pellets were washed by 600 μl methanol two times and air-dried for 5 min, then resuspended in 500 μL SDT-lysis buffer and incubated on Eppendorf Thermomixer for 30 min at l,250 rpm while shaking. The biotin-depleted samples were diluted with PBS buffer (0.1 M phosphate, 0.15 M NaCl, pH 7.2) to a final concentration of 0.5% SDS. Streptavidin-agarose beads (Thermo Fisher Scientific) were washed with PBS buffer three times. After adding 50 μl beads to each sample, the samples were incubated on a rolling wheel at RT for 20 h. The beads were centrifuged for three min at 3500 rpm and removed the supernatant, then washed once with 2 mL PBS buffer containing 2% SDS and six times with 10 mL PBS buffer. An aliquot of the beads was used for immunoblotting, the antibody used was Strep-HRP (1:1,000 dilution; Sigma-Aldrich).

### Identification of proteins by LC-MS

Proteins from HA enrichment were submitted to an on-bead digestion. In brief, dry beads were re-dissolved in 25 μL digestion buffer 1 (50 mM Tris, pH 7.5, 2M urea, 1mM DTT, 5 ng/μL trypsin) and incubated for 30 min at 30 °C in a Thermomixer with 400 rpm. Next, beads were pelleted and the supernatant was transferred to a fresh tube. 50 μL digestion buffer 2 (50 mM Tris, pH 7.5, 2M urea, 5 mM CAA) was added to the beads, after mixing the beads were pelleted, the supernatant was collected and combined with the previous one. The combined supernatants were then incubated o/n at 32 °C in a Thermomixer with 400 rpm; samples were protected from light during incubation. The digestion was stopped by adding 1 μL TFA and desalted with C18 Empore disk membranes according to the StageTip protocol (Rappsilber *et al.*, 2003).

For Streptavidin pulldown double amount of buffers were used (Melkonian *et al.*, 2022). Dried peptides were re-dissolved in 2% ACN, 0.1% TFA (10 μL) and diluted to 0.1 μg/μL for analysis. Samples were analyzed using an EASY-nLC 1200 (Thermo Fisher Scientific) coupled to a Q Exactive Plus mass spectrometer (Thermo Fisher Scientific). Peptides were separated on 16 cm frit-less silica emitters (New Objective, 75 μm inner diameter), packed in-house with reversed-phase ReproSil-Pur C18 AQ 1.9 μm resin. Peptides were loaded on the column and eluted for 115 min using a segmented linear gradient of 5% to 95% solvent B (0 min : 5%B; 0-5 min -> 5%B; 5-65 min -> 20%B; 65-90 min ->35%B; 90-100 min -> 55%; 100-105 min ->95%, 105-115 min ->95%) (solvent A 0% ACN, 0.1% FA; solvent B 80% ACN, 0.1%FA) at a flow rate of 300 nL/min. Mass spectra were acquired in data-dependent acquisition mode with a TOP15 method. MS spectra were acquired in the Orbitrap analyzer with a mass range of 300–1750 m/z at a resolution of 70,000 FWHM and a target value of 3×10^6^ ions. Precursors were selected with an isolation window of 1.3 m/z. HCD fragmentation was performed at a normalized collision energy of 25. MS/MS spectra were acquired with a target value of 10^5^ ions at a resolution of 17,500 FWHM, a maximum injection time (max.) of 55 ms and a fixed first mass of m/z 100. Peptides with a charge of +1, greater than 6, or with unassigned charge state were excluded from fragmentation for MS^2^, dynamic exclusion for 30s prevented repeated selection of precursors.

Raw data were processed using MaxQuant software (version 1.6.3.4, http://www.maxquant.org/) with label-free quantification (LFQ) and iBAQ enabled (Cox and Mann, 2008, Tyanova *et al.*, 2016). MS/MS spectra were searched by the Andromeda search engine against a combined database containing the sequences from Zea mays (Zm-B73-REFERENCE-NAM-5.0) from EnsemblPlants (https://plants.ensembl.org/) and sequences of 248 common contaminant proteins and decoy sequences. Trypsin specificity was required and a maximum of two missed cleavages allowed. Minimal peptide length was set to seven amino acids. Carbamidomethylation of cysteine residues was set as fixed, oxidation of methionine and protein N-terminal acetylation as variable modifications. Peptide-spectrum-matches and proteins were retained if they were below a false discovery rate of 1%.

Statistical analysis of the MaxLFQ values was carried out using Perseus (version 1.5.8.5, http://www.maxquant.org/). Quantified proteins were filtered for reverse hits and hits “identified by site” and MaxLFQ values were log2 transformed. After grouping samples by condition only those proteins were retained for the subsequent analysis that had 2 valid values in one of the conditions. Two-sample t-tests were performed using a permutation-based FDR of 5%. Alternatively, quantified proteins were grouped by condition and only those hits were retained that had 3 valid values in one of the conditions. Missing values were imputed from a normal distribution (1.8 downshift, separately for each column). Volcano plots were generated in Perseus using an FDR of 5% and an S0=1. The Perseus output was exported and further processed using Excel. Relative iBAQ values were calculated per column from MaxQuant output, scaled by a factor of 10^6^ and log10 transformed (Melkonian *et al.*, 2022).

### Psychophysiological interactions (PPIs) and gene ontology (GO) enrichment analysis

PPIs analysis was done by using the STRING database (https://string-db.org/) (Szklarczyk *et al.*, 2021). then further analyzed by Cytoscape software (version 3.9.1, https://cytoscape.org/) with CluePedia (version 1.5.9, https://apps.cytoscape.org/apps/cluepedia) and ClueGo (version 2.5.9, https://apps.cytoscape.org/apps/cluego) (Shannon *et al.*, 2003, Bindea *et al.*, 2009, Bindea *et al.*, 2013, Mlecnik *et al.*, 2019). The GO and KEGG enrichment analysis was done by ShinyGO v0.66 (http://bioinformatics.sdstate.edu/go65/) (Ge *et al.*, 2020).

### Yeast-two hybrid assay

The pGBKT7 and pGADT7 plasmids were co-transformed into yeast strain AH109 (Clontech) using the LiCl-PEG (polyethylene glycol) method (Paiano *et al.*, 2019). Yeast transformants were selected on SD-Leu-Trp plates and grown in liquid SD-Leu-Trp for 16 hours before washing in ddH_2_O. The cell density was adjusted to OD_600_=1.0 and a dilution series was dropped out on SD-Leu-Trp and SD-Leu-Trp-His plates. Plates were incubated for five days at 28°C. At least five independent clones were tested for each combination.

### Agrobacterium-mediated transformation of *Nicotiana benthamiana* leaves

Plasmids of interest were transformed into *Agrobacterium tumefaciens* GV3101 pmp90. Single transformants were confirmed by colony PCR and grown in a liquid dYT medium containing the corresponding antibiotics at 28 °C for 16 hours. Bacterial cells were harvested and resuspended in infiltration buffer (10 mM MES, pH 5.6, 10 mM MgCl_2_, 200 μM acetosyringone) to a final OD_600_=1.0. The different cultures were mixed equally and then infiltrated into the leaves of four-week-old *N. benthamiana* plants.

### Co-immunoprecipitation (Co-IP) and immunoblotting

Leaves transformed with the constructs of interest were harvested at three days post infiltration and frozen in liquid nitrogen. Frozen plant material was ground to a fine powder with 1.5 mL ice-cold protein-lysis buffer (50 mM Tris pH 7.5,150 mM NaCl,10 % glycerol, 2 mM EDTA, 10 mM DTT, 0.5 % IGEPAL, 1 mM PMSF) containing cOmplete™ Protease Inhibitor Cocktail (Roche) for protein extraction. The protein extracts were centrifuged twice, and the resulting supernatant was transferred to a fresh tube as input, an aliquot of the input was used for immunoblotting. 5 μL washed Myc magnetic beads (Chromotek) were added to the protein extracts of each sample and samples were then incubated on a rolling wheel at 4°C. Myc magnetic beads were magnetically separated and washed in 1 mL wash buffer three times. The elution proteins were eluted from the beads with 2× SDS loading buffer (125 mM Tris–HCl pH 6.8, 20% glycerol, 4% SDS, 0.04% bromophenol blue) for immunoblotting.

The input or eluted proteins were separated by SDS-polyacrylamide-gel electrophoresis (PAGE) gels using Gel Electrophoresis (Bio-Rad), and blotted onto polyvinylidene fluoride (PVDF) membranes (Merck) using a Trans-Blot Turbo (Bio-Rad) transfer system. The membranes were blocked in TBST with 3% non-fat milk. The antibodies used were mouse anti-HA primary antibody (1:30,000 dilution; Sigma-Aldrich), anti-mouse lgG secondary antibody (1:10,000 dilution; Thermo Fisher Scientific), rabbit anti-myc primary antibody (1:5,000 dilution; Abcam) and anti-rabbit lgG secondary antibody (1:3,000 dilution; Cell Signaling). The membranes were probed with primary antibody for two hours or overnight and secondary antibody for one hour. TBST washed membranes were detected with the SuperSignal luminol-based chemiluminescent substrate (Thermo Fisher Scientific) by the CCD imaging system (ChemiDoc, Bio-RAD).

### Split-luciferase complementation (split-LUC) assay

Split-LUC assays in *N. benthamiana* were performed as described (Zhou *et al.*, 2018). Agrobacterium-mediated transformation of *N. benthamiana* leaves was performed as described above. For luciferase measurements, leaves were harvested two days post-infiltration and sprayed with 1 mM D-luciferin for 10 min in the dark. The luminescence signals of the leaves from three independent plants were detected by the CCD imaging system (ChemiDoc, Bio-RAD).

### Purification of Umsee1 and Activity-based protein profiling (ABPP)

The pET15b-UmSee1ΔSP plasmid was transformed into Rosetta (DE3) competent cells and recombinant protein expression was initiated by 0.1 mM IPTG treatment at OD_600_= 0.6-0.8. Then cells were incubated at 18°C shaking at 120 rpm for 18 h. Cells were harvested and lysed using a microfluidizer. 6His-UmSee1ΔSP protein was purified using affinity chromatography with Ni-NTA column (Thermo Fisher Scientific) and further purified using size exclusion chromatography with HiLoad Superdex 75 16/600 column (GE-Healthcare) (Li and Sousa, 2012). The ABPP assays were performed as described before (Kolodziejek *et al.*, 2011, Misas-Villamil *et al.*, 2017). In brief, maize leaves were ground and resuspended in 50 mM Tris-HCl buffer (pH 7). The supernatant was pre-incubated for 30 minutes with or without 5 μM (final concentration) recombinant 6His-UmSee1ΔSP or with 20 μM (final concentration) Epoxomicin or DMSO. Then, samples were incubated with 1 μM (final concentration) of the proteasome probe MVB072 for 2 h. Samples were denatured at 95°C in 2x SDS loading buffer and separated on SDS-PAGE gels. The probe was detected using the rhodamine filter (excitation: 532 nm, emission: 580 nm) on a ChemiDoc (Bio-Rad). The gel was stained with SyproRuby (Invitrogen) to determine equal loading according to the manufacturer’s instructions. Quantification of the signals was performed by Image J (Schneider *et al.*, 2012), all the images were transformed into gray-scale and 8-bit type for further analysis. The intensity was normalized to the loading control first, then the final signals were normalized to the epoxomicin control.

### Protein degradation assay

Protein degradation assays were performed as described before (Liu *et al.*, 2010). In brief *N. benthamiana* leaves expressing mCherry-6HA UmSee1ΔSP-mCherry-6HA and ZmSIP3-4myc proteins were harvested three days post-infiltration and frozen in liquid nitrogen. Frozen plant material was ground to a fine powder with 1.0 mL ice-cold lysis buffer (250 mM Tris pH 7.5,150 mM NaCl, 10 % glycerol) for protein extraction. The protein extracts were centrifuged twice at 13,000 x g for 20 min, and the resulting supernatant was transferred to a fresh tube. ZmSIP3-4myc protein extracts were mixed in different proportions with mCherry-6HA or UmSee1ΔSP-mCherry-6HA protein extracts and treated with 100 μM (final concentration) proteasome inhibitor MG132 or DMSO for 45 minutes at 28 °C. The reactions were stopped by boiling samples at 95°C in 2x SDS loading buffer for 10 min, and separated on SDS-PAGE gels. Samples were analyzed by western blot using anti-myc and anti-HA at concentrations specified above. The membrane was stained with Coomassie brilliant blue (CBB) to determine equal protein loading.

### Subcellular localization assay

At three days post-transformation of *N. benthamiana* leaves, fluorescent signals were analyzed using a Leica TCS SP8 confocal laser scanning microscope (Leica) using filters for GFP (excitation 458 nm and emission 470-490 nm) and mCherry (excitation 561 nm and emission 590-603 nm).

## Acknowledgments

We would like to thank Weiliang Zuo for the helpful discussion and suggestions, Ute Meyer for the technical support, Anne Harzen for MS sample processing, Katharina Melkonian for helpful practical advice on the TurboID PL approach and Hermen Overkleeft for proving the MVB072 probe. GD; JCMV and IS acknowledge support from the Cluster of Excellence on Plant Sciences (CEPLAS) funded by the Deutsche Forschungsgemeinschaft (DFG, German Research Foundation) under Germany’s Excellence Strategy – EXC 2048/1 – Project ID: 390686111. This work was funded by the European Research Council under the European Union’s Horizon 2020 research and innovation program (consolidator grant conVIRgens, ID 771035) to GD, the DFG Emmy Noether Programme (SA 4093/1-1) to IMLS and the Daimler and Benz Foundation (to IMLS). HN is supported by the Max Planck Society and is grateful for funding by the DFG (NA 946/1-1).

## Author contributions

W.S., I.M.L.S. J.C.M.V., and G.D. designed the research. W.S., S.C.S., H.N., J.C.M.V., I.M.L.S. and G.D. performed the experiments and data analysis. W.S., J.C.M.V., I.M.L.S. and G.D. wrote the paper with contributions from all the other authors.

## Supporting Information

**Fig S1: UmSee1 interacting proteins enrichment by TurboID-based biotin labelling.**

**Fig S2: Protein-protein interaction (PPIs) analysis of proteins that were specifically identified in UmSee1 samples.**

**Fig S3: Gene Ontology (GO) analysis of proteins that were specifically identified in UmSee1 samples.**

**Fig S4: Identified ZmSIPs (ZmSIP1, ZmSIP2, ZmSIP3).**

**Fig S5: Subcellular localization of ZmSIPs and UmSee1 upon expression in *Nicotiana benthamiana.***

**Fig S6: Y2H assays involving SrSee1 and UhSee1 and the ZmSIPs.**

**Table S1: Primers used in this study.**

**Table S2: Fungal strains used in this study.**

**Table S3: Quantification of infection symptoms on maize seedlings at 12 dpi.**

**Table S4: Mass spectrometry data of UmSee1.**

**Table S5: List of identified UmSee1-interacting proteins.**

