## Supplemental Figures for "Exploiting effector - host interactions using TurboID-based proximity labeling"

### Supporting Figures S1 – S6

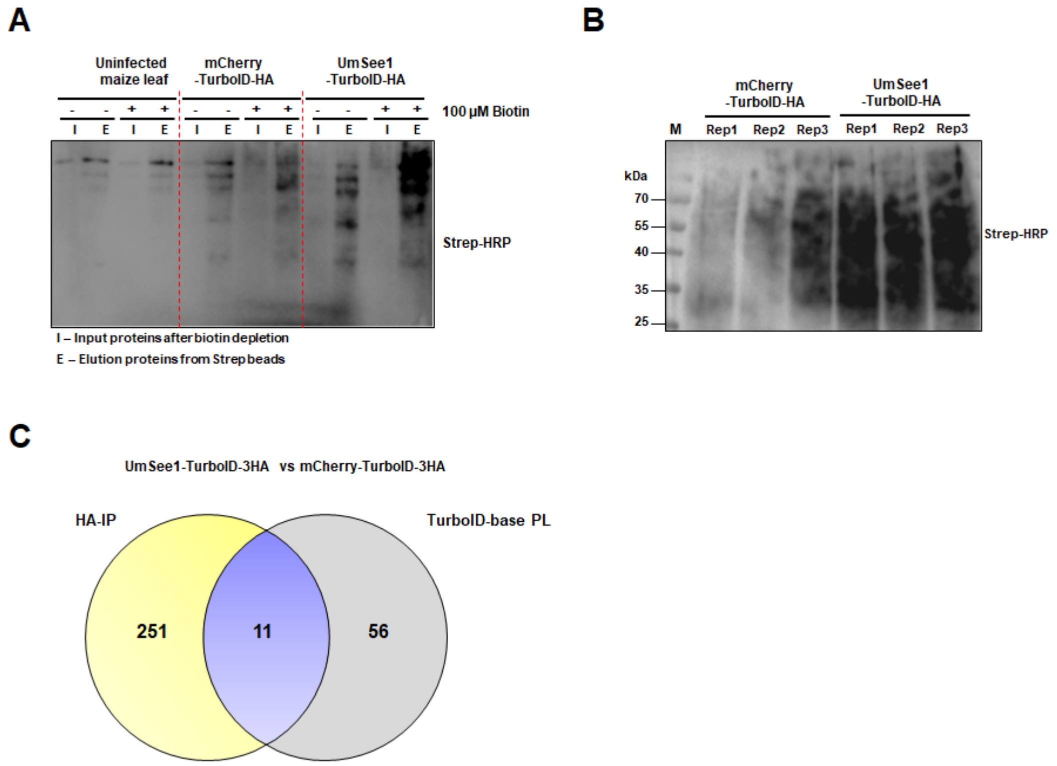


**Fig S1.: UmSee1 interacting proteins enrichment by TurboID-based biotin labelling.** (**A**) Analysis of biotinylated proteins in infected and non-infected leaves. Seven-day-old maize seedlings were infected with *U. maydis* mCherry-TurboID-HA/SG200 or UmSee1-TurboID-HA/ΔSee1. Non-infected leaves were used as a negative control. Two days post-infection non-infected and infected leaves were treated with (+) or without (-) 100 µM biotin. Infected leaves were harvested one-day post biotin treatment for protein extraction and Streptavidin-based precipitation of biotinylated proteins. (**B**) TurboID pull-down of infected *U. maydis* leaves. Biotinylated proteins of three biological replicates (Rep 1-3) were detected using Strep-HRP and further subjected for LC-MS/MS analysis. (**C**) Venn diagrams depicting the proteins that were specifically identified in UmSee1 samples by HA pull-down and TurboID-based biotin labelling.

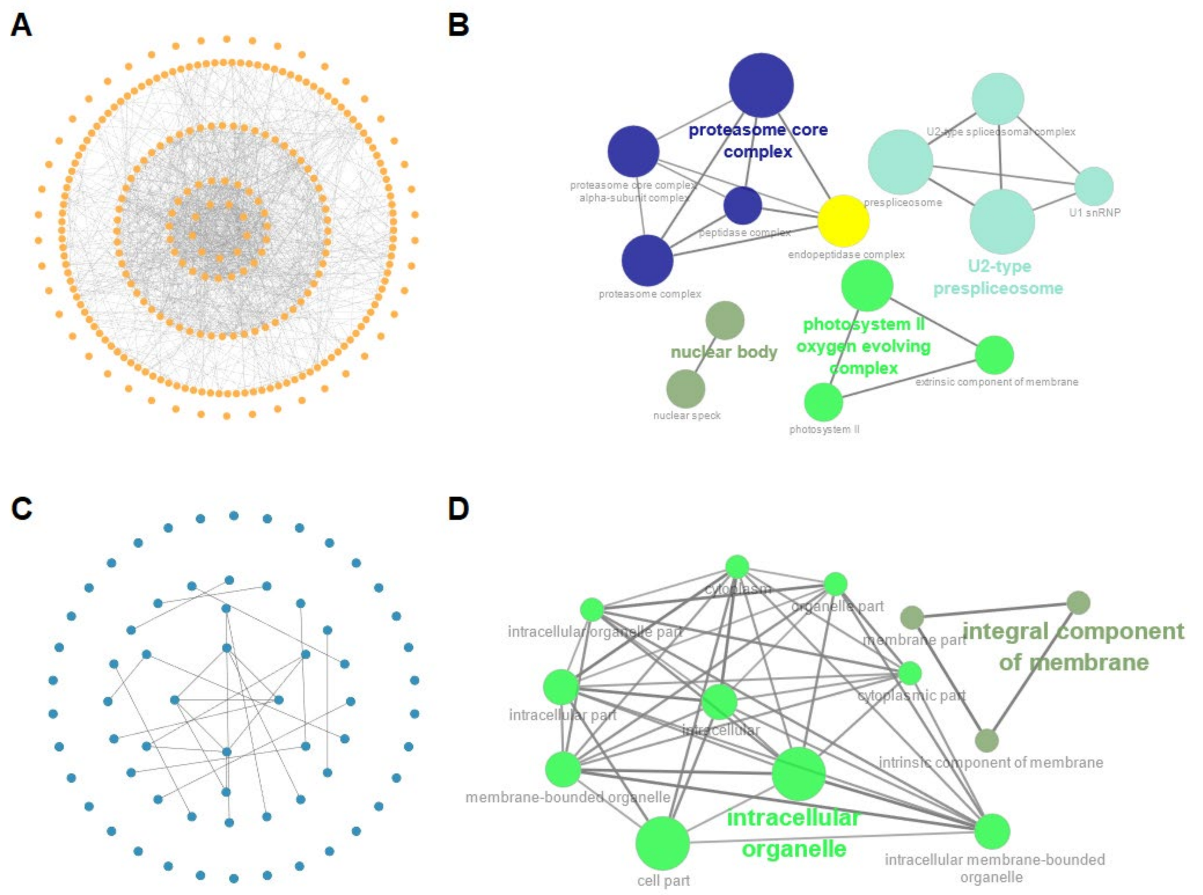


**Fig S2.: Protein-protein interaction (PPIs) analysis of proteins that were specifically identified in UmSee1 samples.** (**A**) Protein-Protein interaction (PPI) analysis of the identified UmSee1 interacting proteins using the STRING database. Significantly enriched UmSee1 interactors from HA-IP were submitted to the STRING database, and the protein interaction network was modified and analyzed by Cytoscape software (version 3.9.1). Edges represent protein-protein associations, including known interactions and predicted interactions (gene neighbourhood, gene fusions, gene co-occurrence). Proteins are represented by their gene ID. (**B**) Cluster analysis of the Protein-Protein interactions network of UmSee1 interactors from the HA-IP dataset by CluePedia (version 1.5.9) and ClueGo (version 2.5.9). (**C**) Protein-Protein interaction (PPI) analysis of the identified UmSee1 interacting proteins using the STRING database. Significantly enriched UmSee1 interactors from TurboID-based PL-IP were submitted to the STRING database, and the protein interaction network was modified and analyzed by Cytoscape software (version 3.9.1). Edges represent protein-protein associations, including known interactions and predicted interactions (gene neighbourhood, gene fusions, gene co-occurrence). Proteins are represented by their gene ID. (**D**) CluePedia (version 1.5.9) and ClueGo (version 2.5.9)- based cluster analysis of the Protein-Protein interactions network of proteins that were specifically identified in UmSee1 samples in the TurboID-based PL dataset.

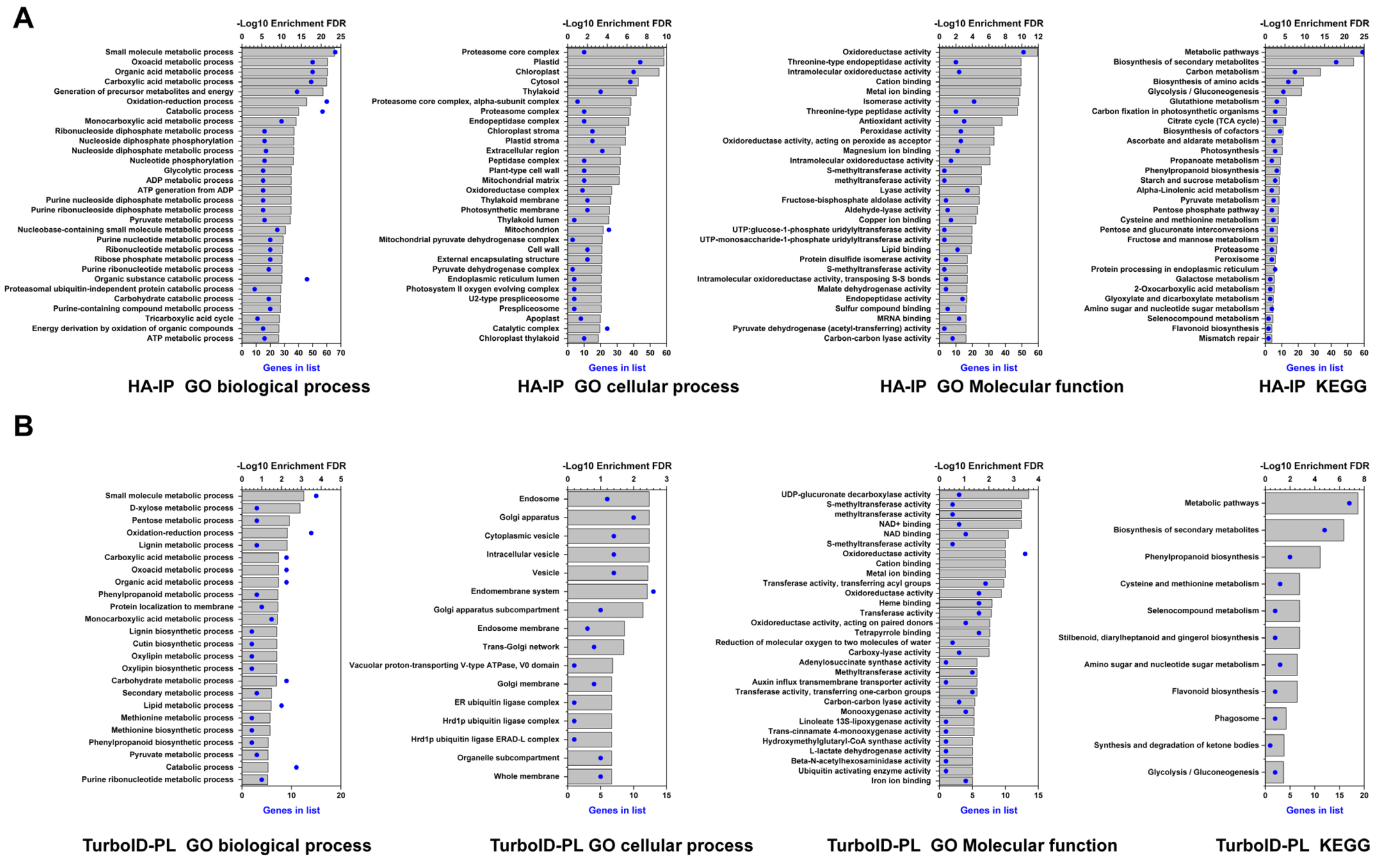


**Fig S3.: Gene Ontology (GO) analysis of proteins that were specifically identified in UmSee1 samples.** (**A**) Gene Ontology (GO) biological, cellular component, molecular function and KEGG enrichment of the identified UmSee1-interacting proteins from the HA-IP dataset using ShinyGO v0.66. (**B**) ShinyGO v0.66-based gene ontology (GO) of biological, cellular component, molecular function and KEGG enrichment of the proteins that were specifically identified in UmSee1 samples in the TurboID-based PL dataset.

**A**

| Name | Protein ID | Funtion |
|---|---|---|
| ZmSIP1 | Zm00001eb157120 | Ubiquitin-conjugating enzyme E2 36 |
| ZmSIP2 | Zm00001eb369210 | Proteasome subunit alpha type 3 |
| ZmSIP3 | Zm00001eb185960 | AAA-type CDC48 protein |

**B**

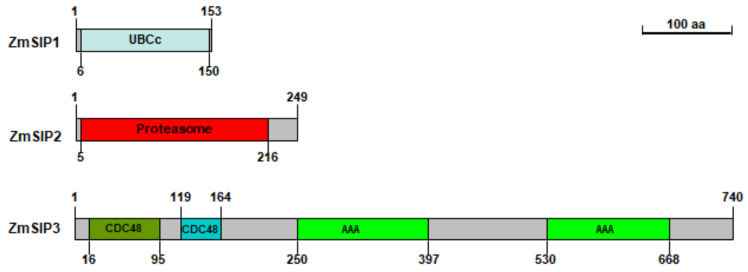


**C**

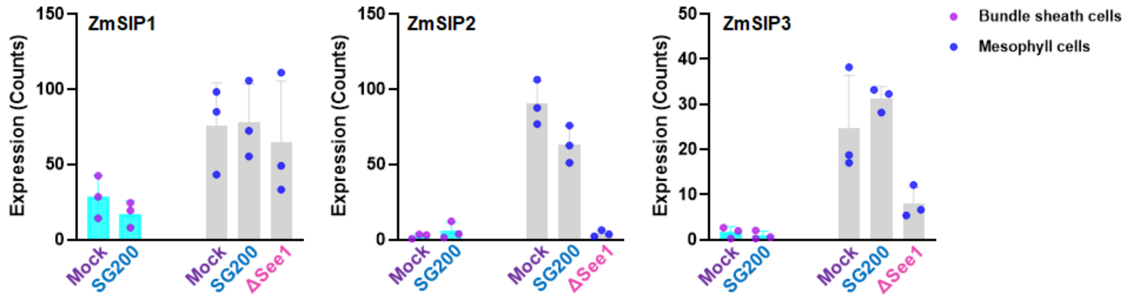


**Fig S4.: Identified ZmSIPs (ZmSIP1, ZmSIP2, ZmSIP3).** (**A**) ZmSIPs protein ID and putative function (**B**) Diagrams that illustrate the protein domain structure of ZmSIP1, ZmSIP2 and, ZmSIP3. (**C**) Expression patterns of the ZmSIP1, ZmSIP2, and ZmSIP3 in different cell type of *U. maydis*-infected maize. Data extracted from the previously published RNA-seq (Villajuana-Bonequi *et al.*, 2019b).

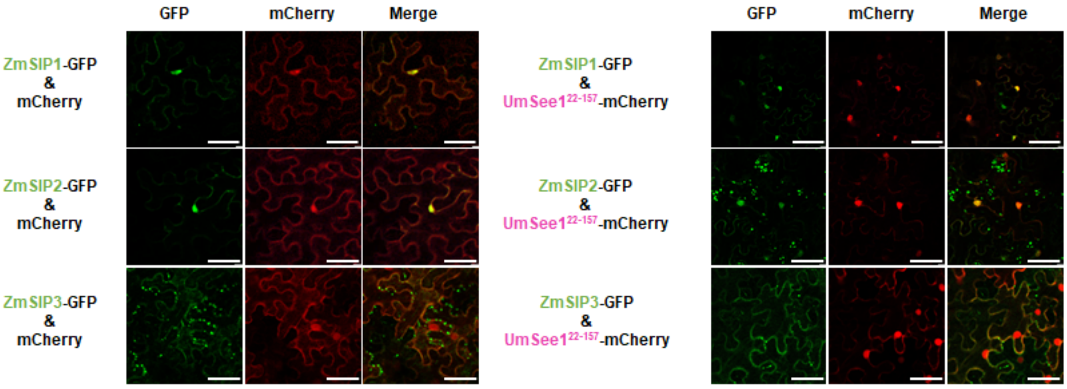


**Fig S5.: Subcellular localization of ZmSIPs and UmSee1 upon expression in *Nicotiana benthamiana*.** UmSee1ΔSP-mCherry, mCherry and ZmSIPs (ZmSIP1, ZmSIP2, ZmSIP3)-GFP were transiently co-expressed in *N. benthamiana* leaves. At three days post-Agrobacterium-mediated transformation fluorescence signals were observed by a Leica TCS SP8 confocal laser scanning microscope (Leica).

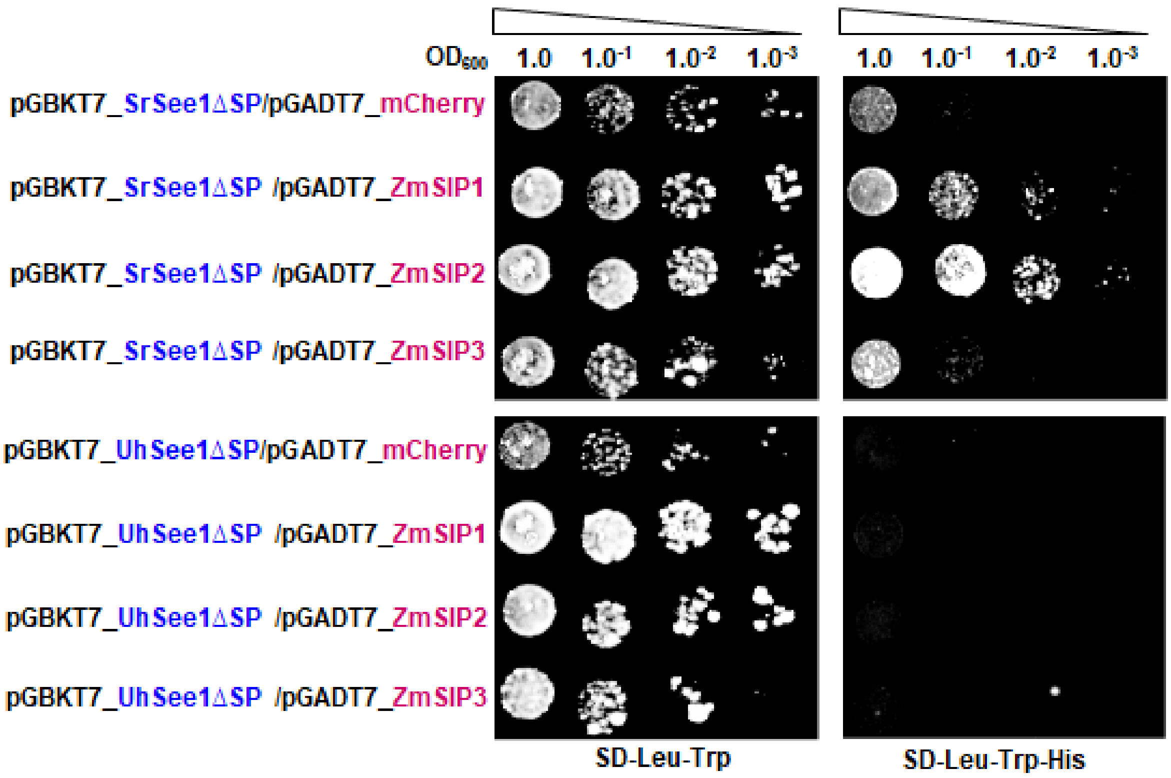


**Fig S6.: Y2H assays involving SrSee1 and UhSee1 and the ZmSIPs.** Yeast strain AH109 was co-transformed with constructs of ZmSIPs (ZmSIP1, ZmSIP2, ZmSIP3) and mCherry, SrSee1 or UhSee1 lacking signal peptide as indicated, and a drop-out series of the double-transformants was grown in the presence (SD-Trp-Leu) and absence (SD-Trp-Leu-His) of histidine. Yeast growth in the absence of histidine suggests the interaction of proteins encoded at the plasmids transformed. Pictures of three independent experiments were taken 5 days after drop out.
