## Supplementary material for "Exploiting effector - host interactions using TurboID-based proximity labeling": Table S1

**Table S1: Primers used in this study.**

| **Primer** | **Sequence (5’-3’)** | **Purpose** |
| --- | --- | --- |
| TurboID-F | cgggcggccgctagctctagaATGAAAGACAATACTGT | ORF of TurboID |
| TurboID-R | ctgcagccgggcggctctagaTTAAGCGTAATCTGGAACAT | ORF of TurboID |
| mCherry-F | cgcagcaccgcggatcccATGGTGAGCAAGGGCGA | ORF of mCherry |
| mCherry-R | tagagctagcggccgcccgCTTGTACAGCTCGTCCA | ORF of mCherry |
| SP_umsee1_-F | cgaccaaacgcagcaccgcggATGCTCTTCACCACCTT | Signal peptide of UmSee1 |
| SP_umsee1_-R | cctcgcccttgctcaccatggGAGCAGACACGTGGACAA | Signal peptide of UmSee1 |
| UmSee1-R | tagagctagcggccgcccCCGTCGTCGGCCCAAATT | ORF of UmSee1 |
| ZmSGT1-F | AGGCCTggatccTCGCGAatgGCCGCGTCGGATCTG | ORF of ZmSGT1 |
| ZmSGT1-R | ATGGTGATGATGACTAGTAATTTCCCACTTCTTGAG | ORF of ZmSGT1 |
| UmSee1-F1 | AGGCCTggatccTCGCGAatgCATCCTCTACAATCG | ORF of UmSee1 |
| UmSee1-R2 | ATGGTGATGATGACTAGTCGTCGTCGGCCCAAATTT | ORF of UmSee1 |
| SrSee1-F | AGGCCTggatccTCGCGAATGCGACCCACCAACACT | ORF of SrSee1 |
| SrSee1-R | ATGGTGATGATGACTAGTCGTGTACGAATCGCCCAC | ORF of SrSee1 |
| UhSee1-F | AGGCCTggatccTCGCGAATGCTCCCACCACGCCAC | ORF of UhSee1 |
| UhSee1-R | ATGGTGATGATGACTAGTACCCCCCTCTCCGCCTCT | ORF of UhSee1 |
| See1-NcoI-F | CATCCATGGCCATGATGCATCCGCTGCAG | ORF of UmSee1 |
| See1-XhoI-R | CATCTCGAGTTAGGTTGTAGGACCAAAC | ORF of UmSee1 |
| ZmSIP1-F | AGGCCTggatccTCGCGAATGGCCAACAGCAACCTC | ORF of ZmSIP1 |
| ZmSIP1-R | ATGGTGATGATGACTAGTTGCACCGCTCGCATACAG | ORF of ZmSIP1 |
| ZmSIP2-F | AGGCCTggatccTCGCGAATGAGCAGCATAGGCACA | ORF of ZmSIP2 |
| ZmSIP2-R | ATGGTGATGATGACTAGTGTCAGCATCCATCTCCTC | ORF of ZmSIP2 |
| ZmSIP3-F | AGGCCTggatccTCGCGAATGGCGGGAAGGTATCAC | ORF of ZmSIP3 |
| ZmSIP3-R | ATGGTGATGATGACTAGTGTAACGGATAATATCACT | ORF of ZmSIP3 |
