## Supplementary material for "Exploiting effector - host interactions using TurboID-based proximity labeling": Table S2

**Table S2: *Ustilago maydis* strains used in this study.**

| **Strain** | **Reference** |
| --- | --- |
| SG200 | Kamper *et al.*, 2006 |
| ΔSee1 | Redkar *et al.*, 2015 |
| mCherry-TurboID-HA/SG200 | This work |
| UmSee1-TurboID-HA/∆See1 | This work |
