## Supplementary material for "Exploiting effector - host interactions using TurboID-based proximity labeling": Table S3

**Table S3 Quantification of infection symptoms on maize seedlings at 12 dpi.**

|  | **Disease Index** | **0** | **1** | **3** | **5** | **7** | **9** | **11** |  |  |  |
| --- | --- | --- | --- | --- | --- | --- | --- | --- | --- | --- | --- |
| **Replicate** | **Strain** | **No symptoms** | **Chlorosis** | **Swelling** | **Small tumors** | **Normal tumors** | **Heavy tumors** | **Dead plants** | **Total plants** | **Total disease index** | **Average disease index** |
| 1 | SG200 | 0 | 0 | 6 | 14 | 32 | 8 | 1 | 61 | **395** | 6,475409836 |
| 1 | mCherry-TurboID-3HA/SG200 | 0 | 0 | 4 | 8 | 26 | 10 | 0 | 48 | **324** | 6,75 |
| 1 | ΔSee1 | 0 | 1 | 2 | 35 | 22 | 2 | 2 | 64 | **376** | 5,875 |
| 1 | UmSee1-TurboID-3HA/ΔSee1 | 0 | 1 | 1 | 9 | 23 | 15 | 1 | 50 | **356** | 7,12 |
| 2 | SG200 | 1 | 2 | 0 | 10 | 28 | 9 | 0 | 50 | **329** | 6,58 |
| 2 | mCherry-TurboID-3HA/SG200 | 0 | 5 | 0 | 5 | 29 | 6 | 4 | 49 | **331** | 6,755102041 |
| 2 | ΔSee1 | 0 | 0 | 5 | 27 | 14 | 4 | 1 | 51 | **295** | 5,784313725 |
| 2 | UmSee1-TurboID-3HA/ΔSee1 | 1 | 1 | 1 | 13 | 19 | 14 | 1 | 50 | **339** | 6,78 |
| 3 | SG200 | 0 | 3 | 0 | 10 | 12 | 8 | 4 | 37 | **253** | 6,837837838 |
| 3 | mCherry-TurboID-3HA/SG200 | 0 | 2 | 1 | 10 | 23 | 6 | 5 | 47 | **325** | 6,914893617 |
| 3 | ΔSee1 | 0 | 6 | 1 | 21 | 8 | 3 | 4 | 43 | **241** | 5,604651163 |
| 3 | UmSee1-TurboID-3HA/ΔSee1 | 0 | 1 | 0 | 11 | 15 | 10 | 2 | 39 | **273** | 7 |

| **Strain** | **No symptoms** | **Chlorosis** | **Swelling** | **Small tumors** | **Normal tumors** | **Heavy tumors** | **Dead plants** | **Total plants** |
| --- | --- | --- | --- | --- | --- | --- | --- | --- |
| **SG200** | 0,68% | 3,38% | 4,05% | 22,97% | 48,65% | 16,89% | 3,38% | **148** |
| **mCherry-TurboID-3HA/SG200** | 0,00% | 4,86% | 3,47% | 15,97% | 54,17% | 15,28% | 6,25% | **144** |
| **ΔSee1** | 0,00% | 4,43% | 5,06% | 52,53% | 27,85% | 5,70% | 4,43% | **158** |
| **UmSee1-TurboID-3HA/ΔSee1** | 0,72% | 2,16% | 1,44% | 23,74% | 41,01% | 28,06% | 2,88% | **139** |

| **Strain** | **disease index 1** | **disease index 2** | **disease index 3** | **Mean** | **Std** |
| --- | --- | --- | --- | --- | --- |
| **SG200** | 6,47541 | 6,58 | 6,837838 | **6,631083** | **0,186536** |
| **mCherry-TurboID-3HA/SG200** | 6,75 | 6,755102 | 6,914894 | **6,806665** | **0,093763** |
| **ΔSee1** | 5,875 | 5,784314 | 5,604651 | **5,754655** | **0,137593** |
| **UmSee1-TurboID-3HA/ΔSee1** | 7,12 | 6,78 | 7 | **6,966667** | **0,172434** |

| **Dunnett's multiple comparisons test** | **Mean Diff.** | **95.00% CI of diff.** | **Below threshold?** | **Summary** | **Adjusted P Value** |
| --- | --- | --- | --- | --- | --- |
| SG200 vs. mCherry-TurboID-3HA/SG200 | -0,1756 | -0.5327 to 0.1815 | No | ns | 0,4036 |
| SG200 vs. ΔSee1 | 0,8764 | 0.5193 to 1.234 | Yes | *** | 0,0003 |
| SG200 vs. UmSee1-TurboID-3HA/ΔSee1 | -0,3356 | -0.6927 to 0.02150 | No | ns | 0,0647 |
